# Hydrogel platform for *in vitro* three-dimensional assembly of human stem cell-derived β cells and endothelial cells

**DOI:** 10.1101/653378

**Authors:** Punn Augsornworawat, Leonardo Velazco-Cruz, Jiwon Song, Jeffrey R. Millman

## Abstract

Differentiation of stem cells into functional replacement cells and tissues is a major goal of the regenerative medicine field. However, one limitation has been organization of differentiated cells into multi-cellular, three-dimensional assemblies. The islets of Langerhans contain many endocrine and non-endocrine cell types, such as insulin-producing β cells and endothelial cells. Transplantation of exogenous islets into diabetic patients can serve as a cell replacement therapy, replacing the need for patients to inject themselves with insulin, but the number of available islets from cadaveric donors is low. We have developed a strategy of assembling human embryonic stem cell-derived β cells with endothelial cells into three-dimensional aggregates on a hydrogel. The resulting islet organoids express β cell markers and are functional, capable of undergoing glucose-stimulated insulin secretion. These results provide a platform for evaluating the effects of the islet tissue microenvironment on human embryonic stem cell-derived β cells and other islet endocrine cells to develop tissue engineered islets.

## 1. Introduction

In diabetes, insulin-producing β cells, which are located within islets of Langerhans in the pancreas, are dysfunctional or destroyed by high levels of metabolites, such as glucotoxicity or lipotoxicity, or by autoimmune attack. The rapid rise in diabetes prevalence has generated much attention towards the development of technologies to better study and treat this disease. However, there is no cure for diabetes, and current treatments are insufficient in controlling the disease for many patients. A small number of patients have been transplanted with cadaveric human islets, which contain β cells, and remained insulin independent for years [1]. Unfortunately this approach is limited because of the scarcity and variability of isolated human islets available for patients, whom often require islets from multiple donors to achieve normal blood sugar levels [2].

Several reports by us and others have detailed approaches for making insulin-producing β-like cells from human embryonic stem cells (hESCs) with the goal that these cells could be used for both cell replacement therapy and drug screening for diabetes [3–8]. These hESC-derived β (SC-β) cells are capable of undergoing glucose-stimulated insulin secretion and express markers found in β cells. While current methodologies produce final cell populations containing SC-β cells, the cellular composition is significantly different than islets found within the body. Current SC-β cell differentiations create cellular clusters that, while visually resembling islets and having functional SC-β cells that are electrically coupled according to Ca^2+^ flux measurements [4], differ significantly from human islets. The differentiation process does produce cells expressing hormones indicative of other islet endocrine cell types, glucagon and somatostatin, but endothelial cells (ECs) are absent. The native islet microenvironment, which is influenced in part by ECs, is highly specialized in supporting islet architecture and function [9]. This three-dimensional islet environment facilitates β cell-β cell and β cell-EC interactions which altogether support β cell survival and insulin secretion. Specifically, ECs produce extracellular matrix (ECM) proteins to provide such interactions [10, 11].

Modulating the microenvironment that SC-β cells experience to be more islet-like is technologically challenging with current approaches. As SC-β cells are derived from endoderm and endothelial cells from mesoderm germ layers, robust simultaneous differentiation of both cell types in the same culture is not possible. This is because current differentiation protocols specify only one germ layer. Approaches focused on assembly of multiple cell types during or after differentiation are therefore preferable. However, currently culture of aggregates containing SC-β cells in suspension is commonly performed on shaker plates or spinner flasks [5], which requires expensive water-, heat-, and CO_2_-resistant equipment and expertise with hESC culture in reactors, or on air-liquid-interfaces [12], which requires manual formation of each individual aggregate and are not amenable to addition of islet components.

Here we established a platform for the assembly of islet like organoids with SC-β cells and ECs. These parameters allow for SC-β cells to assemble and interact with ECs in microtubule networks when cultured on Matrigel. The heterozygous cell assembles express markers found in islets and are capable of undergoing glucose-stimulated insulin secretion. In contrast, SC-β cells and ECs did not assemble when plated in two-dimensional culture on standard tissue culture plastic, in suspension, nor on collagen 1 hydrogels. Such assembly mimics the vascular architect found in native islets, which allows for in vitro based investigations relating to β cell interactions within the microenvironment.

## 2. Materials and methods

### 2.1 Stem cell culture

The HUES8 hESC line was generously provided by Dr. Douglas Melton (Harvard University) and has been previously published [4, 5]. These cells were cultured in an undifferentiated state in mTeSR1 (StemCell Technologies; 05850) in 100-mL or 30-mL spinner flasks (REPROCELL; ABBWVS10A or ABBWVS03A) on a stirrer plate (Chemglass) set at 60 RPM in a humidified incubator set at 5% CO_2_ and 37 °C. Accutase (StemCell Technologies; 07920) was used to passage cells every 3 d. The Vi-Cell XR (Beckman Coulter) was used to quantify viable cell counts and 6 × 10^5^ cells/mL in mTeSR1+ 10 μM Y27632 (Abcam; ab120129) were seeded back into the flasks.

### 2.2 Differentiation to SC-β cells

Differentiation was performed as described by Velazco-Cruz et al [5]. Undifferentiated hESCs were seeded into 30-mL spinner flasks, cultured for 3 d in mTeSR1, and then cultured in the following conditions in order:

<u>Stage 1</u>: 3 days in S1 basal media with 100 ng/ml Activin A (R&D Systems; 338-AC) and 3 μM CHIR99021 (Stemgent; 04-0004-10) for 1 day followed by S1 basal media with 100 ng/ml Activin A for 2 days.
<u>Stage 2</u>: 3 days in S2 basal media with 50 ng/ml KGF (Peprotech; AF-100-19).
<u>Stage 3</u>: 1 day in S3 basal media with 50 ng/ml KGF, 200 nM LDN193189 (Reprocell; 040074), 500 nM PdBU (MilliporeSigma; 524390), 2 μM Retinoic Acid (MilliporeSigma; R2625), 0.25 μM Sant1 (MilliporeSigma; S4572), and 10 μM Y27632.
<u>Stage 4</u>: 5 days in S3 basal media with 5 ng/mL Activin A, 50 ng/mL KGF, 0.1 μM Retinoic Acid, 0.25 μM SANT1, and 10 μM Y27632.
<u>Stage 5</u>: 7 days in S5 basal media with 10 μM ALK5i II (Enzo Life Sciences; ALX-270-445-M005), 20 ng/mL Betacellulin (R&D Systems; 261-CE-050), 0.1 μM Retinoic Acid, 0.25 μM SANT1, 1 μM T3 (Biosciences; 64245), and 1 μM XXI (MilliporeSigma; 595790). At the end of this stage, clusters were reaggregated by dispersion with TrypLE Express (ThermoFisher; 12604013) and replating in a 6-well plate on an Orbi-Shaker (Benchmark).
<u>Stage 6</u>: 12-22 days in enhanced serum-free media (ESFM).

The basal media formulations are as follows:

<u>S1 basal media</u>: 500 mL MCDB 131 (Cellgro; 15-100-CV) plus 0.22 g glucose (MilliporeSigma; G7528), 1.23 g sodium bicarbonate (MilliporeSigma; S3817), 10 g bovine serum albumin (BSA) (Proliant; 68700), 10 μL ITS-X (Invitrogen; 51500056), 5 mL GlutaMAX (Invitrogen; 35050079), 22 mg vitamin C (MilliporeSigma; A4544), and 5 mL penicillin/streptomycin (P/S) solution (Cellgro; 30-002-CI).
<u>S2 basal media</u>: 500 mL MCDB 131 plus 0.22 g glucose, 0.615 g sodium bicarbonate, 10 g BSA, 10 μL ITS-X, 5 mL GlutaMAX, 22 mg vitamin C, and 5 mL P/S.
<u>S3 basal media</u>: 500 mL MCDB 131 plus 0.22 g glucose, 0.615 g sodium bicarbonate, 10 g BSA, 2.5 mL ITS-X, 5 mL GlutaMAX, 22 mg vitamin C, and 5 mL P/S.
<u>S5 media</u>: 500 mL MCDB 131 plus 1.8 g glucose, 0.877 g sodium bicarbonate, 10 g BSA, 2.5 mL ITS-X, 5 mL GlutaMAX, 22 mg vitamin C, 5 mL P/S, and 5 mg heparin (MilliporeSigma; A4544).
<u>ESFM</u>: 500 mL MCDB 131 plus 0.23 g glucose, 10.5 g BSA, 5.2 mL GlutaMAX, 5.2 mL P/S, 5 mg heparin, 5.2 mL MEM nonessential amino acids (Corning; 20-025-CI), 84 μg ZnSO_4_ (MilliporeSigma; 10883), 523 μL Trace Elements A (Corning; 25-021-CI), and 523 μL Trace Elements B (Corning; 25-022-CI).

### 2.3 Light microscopy

Images of cell clusters stained with 2.5 μg/mL DTZ (MilliporeSigma; 194832) were taken with an inverted light microscope (Leica DMi1).

### 2.4 Immunohistochemistry

Clusters were fixed with 4% paraformaldehyde (Electron Microscopy Science; 15714) overnight at 4 °C, embedded in Histogel (Thermo Scientific; hg-4000-012), and paraffin-embedded and sectioned by the Division of Comparative Medicine (DCM) Research Animal Diagnostic Laboratory Core at Washington University. Immunostaining was performed by paraffin removal with Histoclear (Thermo Scientific; C78-2-G), rehydration by treatment with increasing ratios of water to ethanol, antigens retrieved by treatmetn with 0.05 M EDTA (Ambion; AM9261) in a pressure cooker (Proteogenix; 2100 Retriever). Non-specific antibody binding was blocked with a 30-min treatment in staining buffer (5% donkey serum (Jackson Immunoresearch; 017-000-121) and 0.1% Triton-X 100 (Acros Organics; 327371000) in PBS), followed by overnight staining with 1:300 dilutions of rat-anti-C-peptide (DSHB; GN-ID4-S) and mouse-anti-glucagon (ABCAM; ab82270) primary antibodies. Samples were stained with donkey secondary antibodies containing Alexa Fluor fluorophores (Invitrogen) for 2 hr at 4 °C, and treated with DAPI in the mounting solution Fluoromount-G (SouthernBiotech; 0100-20). Imaging was performed on a Nikon A1Rsi confocal microscope.

### 2.5 Assembly of SC-β cells and ECs in suspension

Human umbilical vein endothelial cells (HUVECs) were purchased form Lonza (C2519A) and cultured in EGM-2 media (Lonza; CC-3162). HUVECs and Stage 6 clusters containing SC-β cells were dispersed with TripLE Express and plated into a 6-well plate on an Orbi-Shaker at 100 rpm. Two conditions were tested: 3×10^6^ Stage 6 cells only (control) and 2.5×10^6^ Stage 6 cells mixed with 0.5×10^6^ HUVECs. Cells were cultured in a media of 90% ESFM and 10% EGM-2. The presence of these cell types was assessed after 48 hr by dispersion and plating into a 96-well plate followed by immunostaining.

### 2.6 Whole-mount immunostaining

Cell assemblies were fixed within the well using 4% paraformaldehyde treatment overnight at 4 °C. Non-specific antibody binding was blocked with a 30-min treatment in staining buffer (5% donkey serum (Jackson Immunoresearch; 017-000-121) and 0.1% Triton-X 100 (Acros Organics; 327371000) in PBS) followed by overnight staining with 1:300 dilutions of rat-anti-C-peptide (DSHB; GN-ID4-S) and mouse-anti-CD31 (Dako; M082329-2) primary antibodies. Samples were stained with donkey secondary antibodies containing Alexa Fluor fluorophores (Invitrogen) for 2 hr at 4 °C and treated with DAPI. Imaging was performed on a Nikon A1Rsi confocal microscope.

### 2.7 Assembly of SC-β cells and ECs in two-dimensional culture

HUVECs and Stage 6 clusters containing SC-β cells were dispersed with TripLE Express and plated (1×10^5^ cells each) into a 96-well plate. Cells were cultured in a media of 90% ESFM and 10% EGM-2. After 24 hr, the cells were fixed with 4% paraformaldehyde for immunostaining assessment of both cell types.

### 2.8 Assembly of SC-β cells and ECs on collagen 1 and Matrigel

Cold Matrigel (Fisher; 356230; 60 μL) was transferred into 96-well plates and then incubated at 37 °C for 1 hr to create gel slabs. Collagen 1 (ThermoFisher; A1048301; 60 μL) at 4 mg/mL pH neutralized, transferred into 96-well plates, and incubated at RT for 1 min to create gel slabs. After this time, HUVECs and Stage 6 clusters containing SC-β cells were dispersed with TripLE Express and plated on top of the gel slabs over a range of cell numbers (0.5-2×10^5^ Stage 6 and 0-1.5×10^5^ HUVECs). Cells were cultured in a media of 90% ESFM and 10% EGM-2. After 24 hr, the cells were fixed with 4% paraformaldehyde for immunostaining assessment of both cell types.

### 2.9 Glucose-stimulated insulin secretion assay

This assay was performed by first washing cells twice with KRB buffer (128 mM NaCl, 5 mM KCl, 2.7 mM CaCl_2_ 1.2 mM MgSO_4_, 1 mM Na_2_HPO_4_, 1.2 mM KH_2_PO_4_, 5 mM NaHCO_3_, 10 mM HEPES (Gibco; 15630-080), and 0.1% BSA) and equilibrating cells in 2 mM glucose KRB for a 1 hr. The supernatant was removed and replaced with fresh 2 mM glucose KRB, discarding the old KRB solution. Assemblies were incubated for 1 hr, the supernatant removed and replaced with fresh 20 mM glucose KRB, retaining the old KRB solution. The assemblies were incubated for 1 hr, the supernatant removed and retained. Insulin concentration with the retained KRB supernatant was quantified with a Human Insulin ELISA (ALPCO; 80-INSHU-E10.1). Secretion was normalized to cell counts by single-cell dispersing assemblies with 10-min TrypLE Express treatment and quantifying viable cell count with a Vi-Cell XR.

### 2.10 Real-time PCR

The RNeasy Mini Kit (Qiagen; 74016) with DNase treatment (Qiagen; 79254), was used to extract RNA from cell assemblies. The High Capacity cDNA Reverse Transcriptase Kit (Applied Biosystems; 4368814) was used to make cDNA for gene expression measurements. Real-time PCR was performed with a StepOnePlus (Applied Biosystems) instrument using PowerUp SYBR Green Master Mix (Applied Biosystems; A25741). Analysis was performed using ∆∆Ct methodology and normalization to TBP. The following primer pairs used: CAATGCCACGCTTCTGC, TTCTACACACCCAAGACCCG; <u>PDX1</u>, CGTCCGCTTGTTCTCCTC, CCTTTCCCATGGATGAAGTC; <u>TBP</u>, GCCATAAGGCATCATTGGAC, AACAACAGCCTGCCACCTTA; <u>NKX6-1</u>, CCGAGTCCTGCTTCTTCTTG, ATTCGTTGGGGATGACAGAG; <u>CHGA</u>, TGACCTCAACGATGCATTTC, CTGTCCTGGCTCTTCTGCTC; <u>NEUROD1</u>, ATGCCCGGAACTTTTTCTTT, CATAGAGAACGTGGCAGCAA; <u>NKX2-2</u>, GGAGCTTGAGTCCTGAGGG, TCTACGACAGCAGCGACAAC; <u>GCK</u>, ATGCTGGACGACAGAGCC, CCTTCTTCAGGTCCTCCTCC; <u>MAFB</u>, CATAGAGAACGTGGCAGCAA, ATGCCCGGAACTTTTTCTTT.

### 2.11 Statistical analysis

GraphPad Prism was used to determine statistical significance using two-sided unpaired and paired *t*-tests. Data is represented as mean±SEM.

## 3. Results

### 3.1 Development of platform for SC-β cells and EC assembly

We have previously published a protocol for the generation of SC-β cells from hESCs [5] (**Fig. 1A**). This approach is done entirely in suspension culture with the cells grown as aggregates (**Fig. 1B-C**). Specific growth factors and molecules are given in different combinations as differentiation progresses to recapitulate pancreatic development. At the end of the protocol, we generate populations of cells that expression C-peptide and other islet hormones, including glucagon (**Fig. 1D**).

**Fig. 1.**
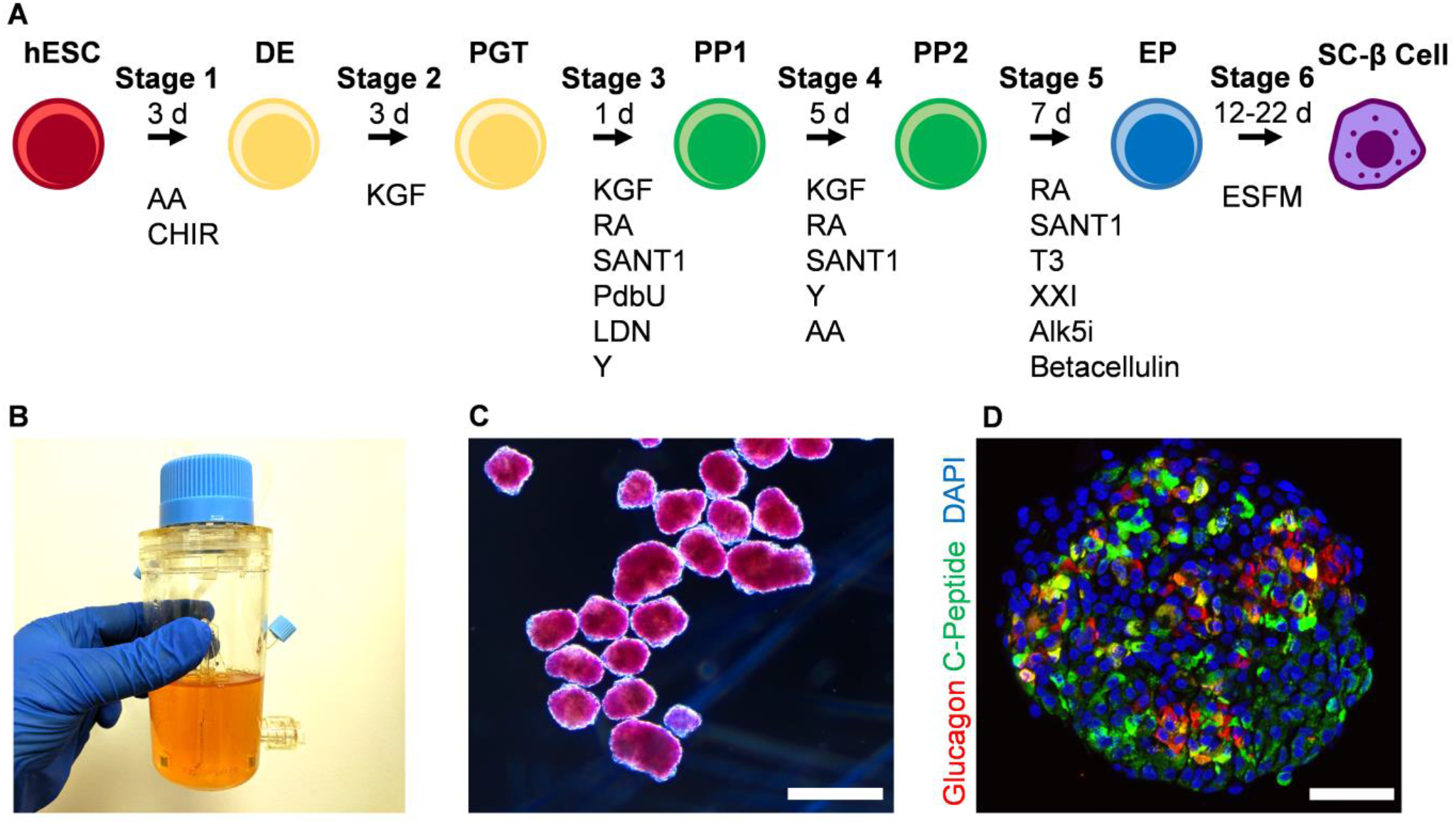
Generation of SC-β cells. **A**. Schematic diagram of differentiation process to generate SC-β cells. **B**. Image of spinner flask approach used to grow and differentiate hESCs to SC-β cells. **C**. Micrograph of Stage 6 clusters stained with dithizone, a dye that stains β cells red, imaged under bright field. Scale bar = 500 μm. **D**. Immunostaining of Stage 6 cluster sectioned and stained for C-peptide and glucagon. Scale bar = 100 μm. DE, definitive endoderm; PGT, primitive gut tube; PP1, pancreatic progenitor 1; PP2, pancreatic progenitor 2; EP, endocrine progenitor; AA, activin A; CHIR, CHIR9901; KGF, keratinocyte growth factor; RA, retinoic acid; Y, Y27632; LDN, LDN193189; PdbU, phorbol 12,13-dibutyrate; T3, triiodothyronine; Alk5i, Alk5 inhibitor type II; ESFM, enriched serum-free medium.

While this differentiation protocol produces SC-β cells, ECs are absent, in contrast to what is seen in native human islets. In order to develop a platform that enables study of SC-β cells and ECs, we first attempted to disperse the SC-β cell clusters our protocol normally generates, mix with a single cell dispersion of ECs, and allow them to spontaneously reaggregate in a 6-well plate on an orbital shaker at 100 rpm, as we have used to previously reaggregated SC-β cell clusters [5] (**Fig. 2**). The morphology of the resulting clusters was unaffected by the inclusion of ECs (**Fig. 2A**). To check for the incorporation of ECs, we dispersed and plated the reaggregated clusters, then stained for C-peptide, a β cell marker produced by the insulin gene, and CD31, an endothelial cell marker (**Fig. 2B**). We observed little to no CD31+ cells, indicating this approach did not enable ECs to be incorporated with the SC-β cell clusters. In a separate experiment, we plated a single-cell dispersion of SC-β cells mixed with ECs on the bottom of a multi-well plate and assessed with immunostaining (**Fig. 3**). While we were able to observe many C-peptide+ and CD31+ cells, these populations tended to segregate away from each other, with physical proximity of the populations being observed only in rare instances.

**Fig. 2.**
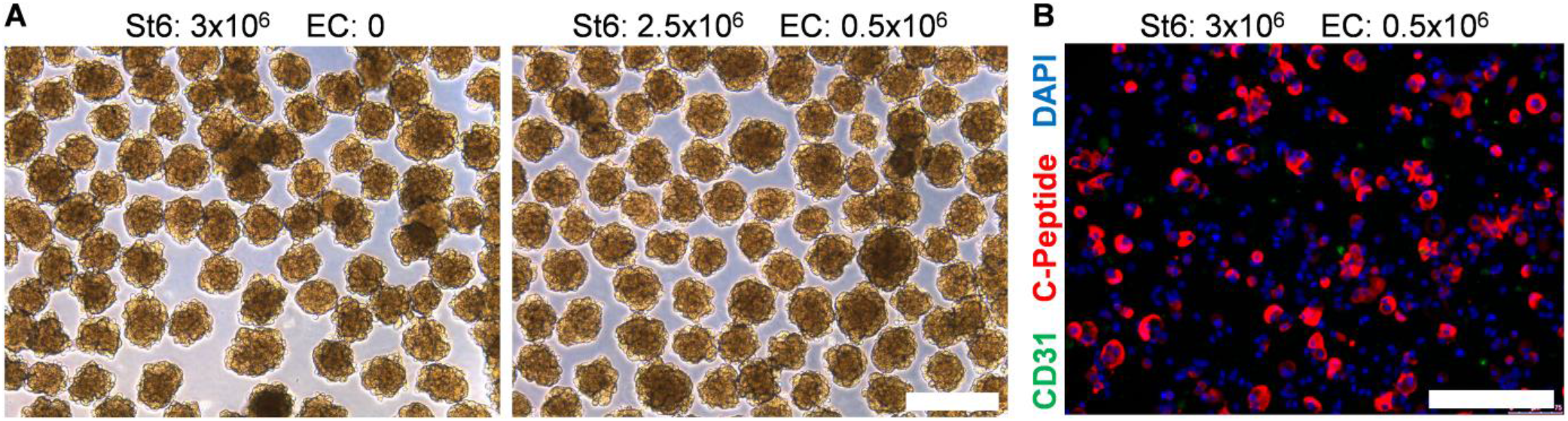
ECs and SC-β cells do not assemble in suspension culture. **A**. Micrographs of unstained reaggregated Stage 6 clusters with or without the addition of ECs. Scale bar = 400 μm. **B**. Immunostaining of Stage 6 clusters reaggregated with ECs dispersed and plated for assessment. Scale bar = 150 μm.

**Fig. 3.**
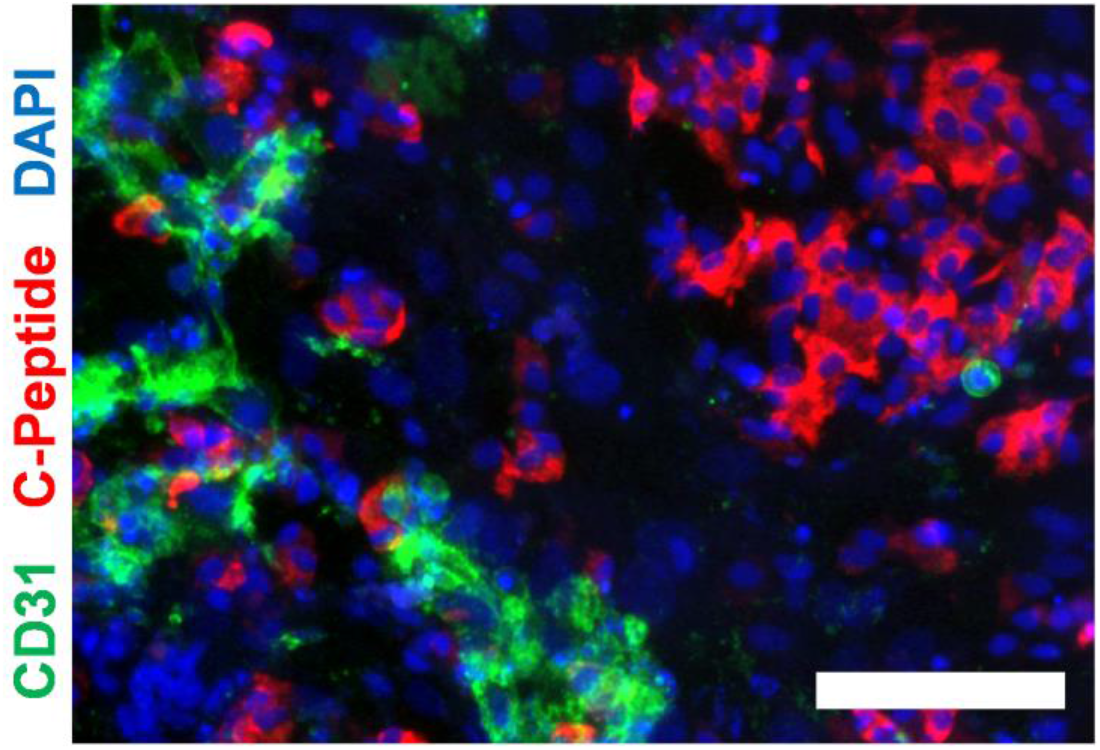
ECs and SC-β cells do not assemble in two-dimensional culture. Immunostaining of Stage 6 clusters mixed with ECs and plated for assembly. Scale bar = 150 μm.

### 3.2 Hydrogel platform enables *SC-β cells and EC assembly*

After observing the difficulty of facilitating C-peptide+ and CD31+ cell physical association, we turned to assembly on top of hydrogels (**Fig. 4**). Two common commercially available hydrogels were assessed: Rat tail-derived collagen 1 and Matrigel, which is a protein mixture derived from mouse sarcoma cells that consists in part of basement membrane extracellular matrix proteins. After creating slabs of each of the polymerized hydrogels, a mixture of single-cell dispersed SC-β cells and ECs at varying ratios was dispensed on top, and assembly of cells observed after 24 hr. At all cell ratios tested, no notable cell aggregates were observed with collagen 1. However, on Matrigel, both 1:1 and 3:1 ratios of SC-β cell to EC produced three-dimensional structures reminiscent of microtubule networks [13]. Higher ratios of ECs tended to produce more sheet-like morphologies, similar to what was observed with collagen I. SC-β cells without endothelial cells produced small aggregates, which is interesting because this did not require the normal equipment used for SC-β cell culture and aggregation: Stirrer, shakers, and/or spinner flasks. Taken together, these data demonstrate that three-dimensional assembly of SC-β cells and ECs can be achieved using polymerized Matrigel with a cell ratio of 1:1 to 3:1.

**Fig. 4.**
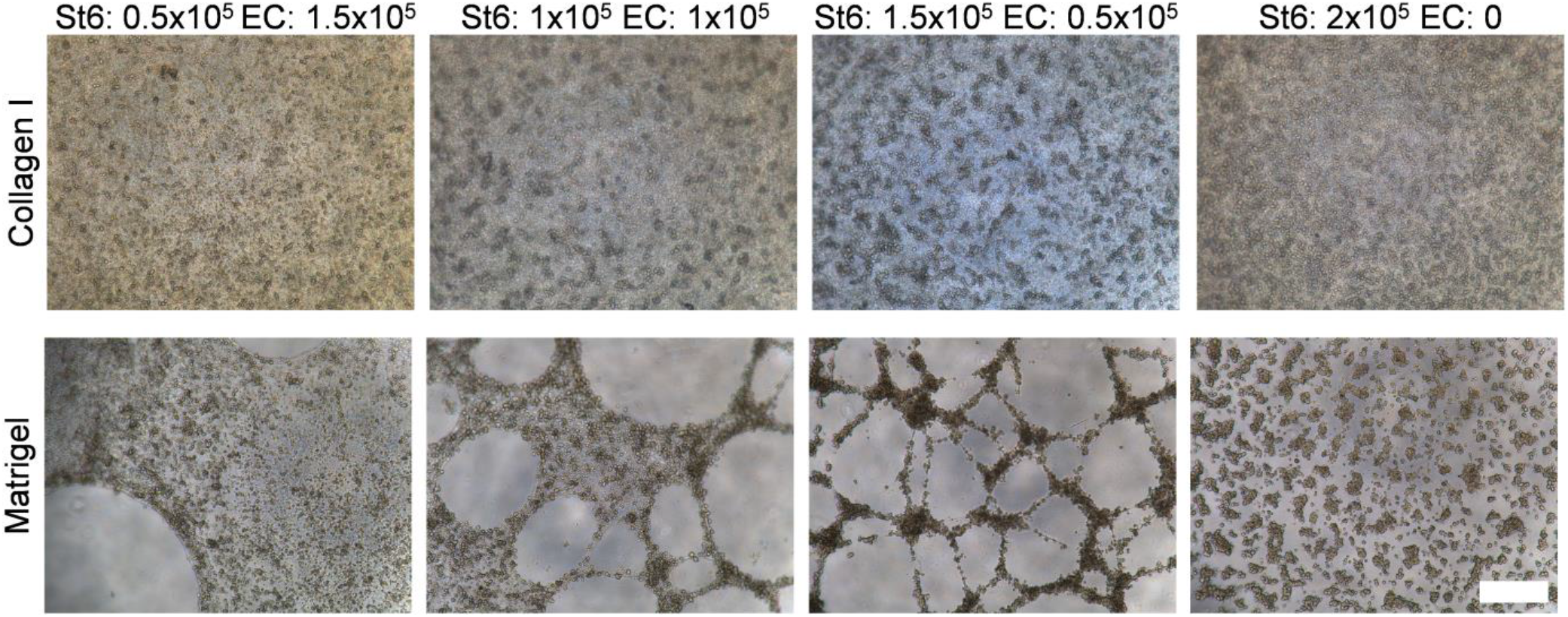
ECs and SC-β cells assemble on top of Matrigel but not collagen 1 gels. Shown are micrographs of varying ratios of ECs and SC-β cells after 24 hr. Scale bar = 400 μm.

### 3.3 Characterization of islet organoids

To characterize islet organoid assembly using our developed platform, we whole-mount stained the resulting aggregate and imaged it with confocal microscopy (**Fig. 5**). We confirmed that the microtubule was formed by the ECs (CD31+) cells. We also observed that the C-peptide+ cells were mostly present on top of the resulting microtubule network, with a few non-assembled C-peptide+ cells observed. Based on our prior observation of the dispersed nature of SC-β cell clusters on Matrigel in the absence of ECs (**Fig. 4**), it is likely that the ECs are secreting pro-migratory factors that attract SC-β cells to the microtubule network.

**Fig. 5.**
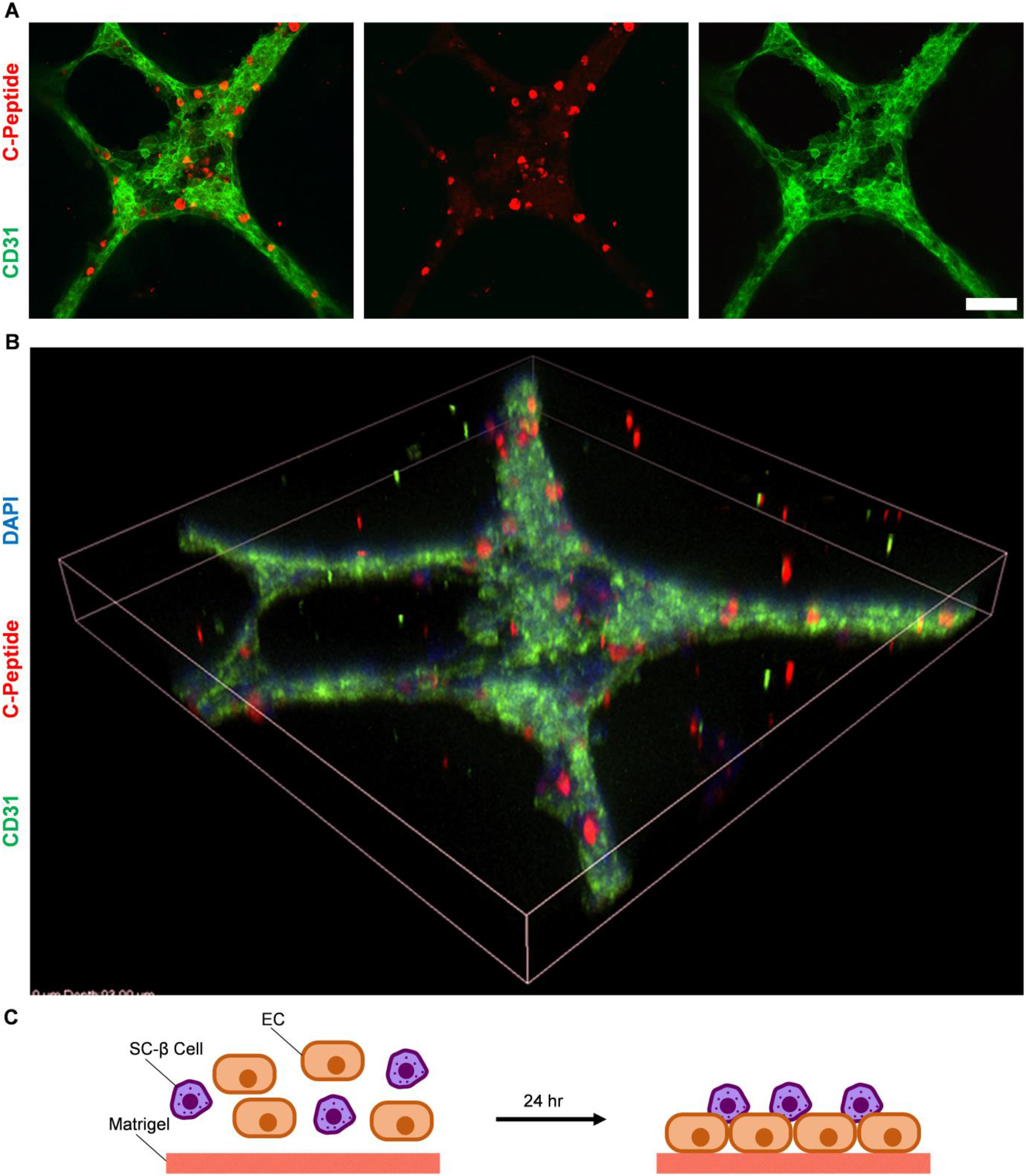
Immunostaining assessment of ECs and SC-β cells assemble formed on Matrigel. **A**. En face image of immunostained organoid. Scale bar = 100 μm. **B**. Tilted view of constructed three-dimensional z-stacks. The shown dimensions are 636.4 × 636.4 × 93 μm. **C**. Schematic cartoon of islet organoid assembly.

To evaluate the potential of this platform for islet organoid assembly for evaluating islet microenvironment parameters, we performed a glucose-stimulated insulin secretion assay of SC-β cell/EC assemblies (**Fig. 6**). The normal physiological function of pancreatic β cells in the body is to secrete insulin in response to high glucose stimulation. This *in vitro* assay involves treating cells first with low (2 mM) glucose for an hour, collecting the resulting supernatant, then subsequently treating cells with high (20 mM) glucose for an hour, collecting the resulting supernatant, and quantifying the amount of insulin released with ELISA. Testing three independent replicates of the islet organoid assembly revealed all three were robustly functional by secreting higher insulin at high glucose (p<0.01; two-way paired *t*-test). On average, insulin secretion increased by 3.9±0.3x by high glucose stimulation. These data show that the assembled islet organoids are glucose-responsive and secrete insulin.

**Fig. 6.**
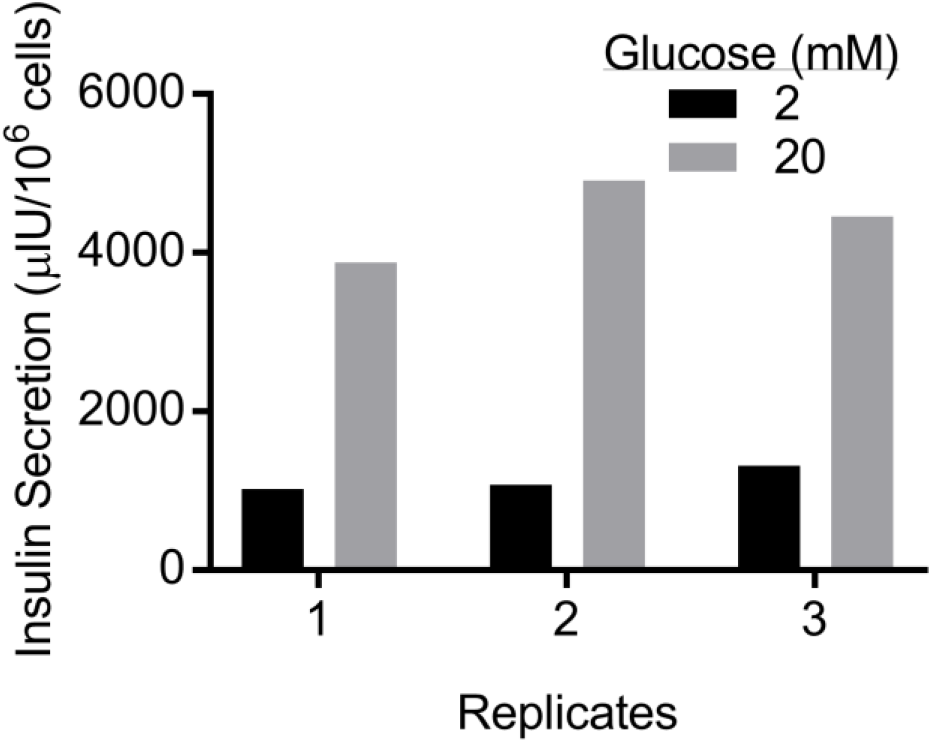
Glucose-stimulated insulin secretion assay of EC and SC-β cell assembly formed on Matrigel. Three biological replicates are plotted separately.

Finally, we evaluated our SC-β cell/EC assemblies by expression of β cell genes (**Fig. 7**). Key genes associated with β cell identity, including insulin (INS), transcription factors (MAFB, PDX1, NKX6-1, NKX2-2, NEUROD1), and with function (CHGA, GCK) were all highly expressed in our assembled islet organoids compared to undifferentiated hESC controls. These data show that SC-β cell assemblies made with our platform express β cell markers.

**Fig. 7.**
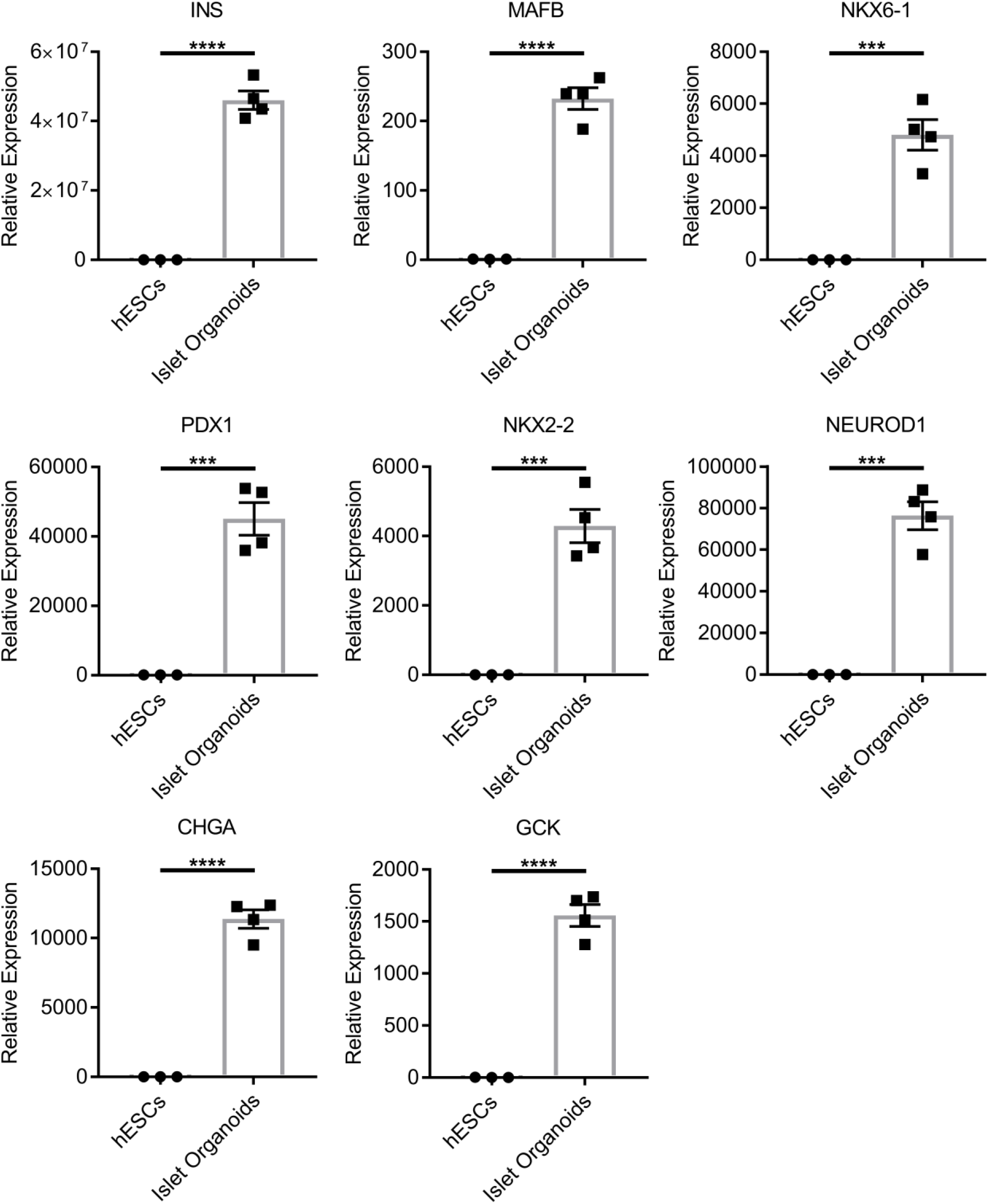
Gene expression analysis of EC and SC-β cell assembly formed on Matrigel. Eight genes associated with the β cells were measured and comparing the islet organoid (n=4) to undifferentiated hESCs (n=3). ***p<0.001 and ****p<0.0001 by two-way unpaired *t*-test.

## 4. Discussion

The islets of Langerhans are complex, multicellular tissues that are responsible for maintaining glucose tolerance within humans through their ability to sense glucose and secrete hormones. Islets consist of β cells and other endocrine cell types along with ECs. Here we developed a platform that enables the assembly of SC-β cells with ECs. The assembled heterogenous cell mixture were capable of undergoing glucose-stimulated insulin secretion, a key β cell functional feature, and expressed a panel of β cell genes that are associated with its identity and function. We encountered difficulties successfully assembling SC-β cells with ECs using other approaches, indicating that the conditions for this phenotype are limited. Specifically, we found the plating on top of hydrogels made from Matrigel and a SC-β cell to EC ratio of 3:1 to be optimal, with SC-β cells sitting on top of microtubule networks formed by the ECs.

Current protocols produce SC-β cells that resemble their *in vivo* counterparts in many important parameters but are still different in many important aspects. While SC-β cells are capable of undergoing glucose-stimulated insulin secretion and controlling blood sugar levels in mice, how glucose-responsive the cells are and how much insulin the cells secrete is still lower than primary cadaveric human islets [4–6, 12, 14]. SC-β cells express many markers found in primary cadaveric human islets, but several genes associated with maturation continue to be under expressed, such as MAFA and UCN3 [5, 14, 15]. We hope our reported platform for assembling key islet components enables future studies to identify the parameters to make more mature SC-β cells in tissue engineered islets. A practical feature of our platform is that it can be achieved without specialized reagents, equipment, or training, which will facilitate future studies and is in contrast with current SC-β cell culture methodologies [5, 12].

Approaches have been developed in order to generate SC-β cell-containing aggregates that better resemble native islets. Optimization of the timing and combinations of soluble small molecules and growth factors have led to increases in differentiation yield [5]. Resizing of clusters has led to increased function both *in vitro* and *in vivo* [5, 6, 14, 16], in part by limiting hypoxia [16]. Sorting based on a transgenic reporter [6] or surface marker [14] has allowed further increases in the purity of SC-β cells to better define the cellular population present in aggregates. Combining 3D printing with fibrin gels has better enabled transplantation of resized SC-β cell clusters [16]. Addressing these differences and inefficiencies will bring stem cell technology closer to clinical translation and increase our understanding of β cell biology and diabetes pathology.

While we did not find success with several approaches that we attempted, alternative methodologies could be developed to assemble ECs and SC-β cells. Candiello et al. reported a system of combining ECs with hESCs differentiated to insulin-producing cells [17] but only achieved insulin secretion per cell that was order 10^2^ lower than achieved with our platform here. We recently used microwells to resize and enable embedding of SC-β cell aggregates into fibrin gels within a 3D-printed macroporous device before transplantation, but introduction of ECs was not studied [16].

Tissue engineered islet organoids have value in diabetes cell replacement therapy and drug screening. Currently a major challenge in the field of diabetes cell replacement therapy is sourcing of sufficient numbers of glucose-responsive insulin-secreting β cells [18]. Differentiated hESCs offer a potentially unlimited number of cells for this purpose [19], and this approach is enhanced by improving the maturation of the generated SC-β cells [20]. The inclusion of endothelial cells with SC-β cells could be combined with macroporous scaffolds that enable retrievability [16, 21] or other beneficial materials [22–26] to develop a more comprehensive transplantation strategy for diabetes. An alternative cell type to SC-β cells are earlier pancreatic progenitors derived from hESCs. These are currently being explored for cell replacement therapy, as these cells can spontaneously differentiate into SC-β cells after transplantation over the course of months [3, 27–29], but the mechanism of this maturation process is unknown and differs based on rodent species [27]. SC-β cells or earlier progenitors, particularly derived from diabetic patients through an induced pluripotent stem cell (iPSC) intermediate, are currently being studied for disease modeling and drug screening purposes [3, 30–41]. These studies would benefit from a cellular assembly that more closely reflects the native islet microenvironment [42].

## 5. Conclusions

Our findings show that assembly of SC-β cells with ECs will only occur under specific culture conditions. Spontaneous aggregation into clusters, assembly on two-dimensional convention tissue culture plastic, and plating on top of collagen 1 gels were unable to achieve assembly. However, we observed that dispersing and plating SC-β cells with ECs in a 3:1 ratio on-top of Matrigel provided the greatest assembly. These assembled islet organoids were able to undergo glucose-stimulated insulin secretion and expressed a panel of β cell markers. Our described approach provides a platform the study of key microenvironmental components for development of tissue engineered islet for diabetes cell replacement therapy and drug screening.

## Acknowledgements

This work was supported by the NIH (R01DK114233), JDRF Career Development Award (5-CDA-2017-391-A-N), Washington University Center of Regenerative Medicine, and startup funds from Washington University School of Medicine Department of Medicine. L.V.C. was supported by the NIH (R25GM103757). Microscopy was performed through the Washington University Center for Cellular Imaging (WUCCI), which is supported by Washington University School of Medicine, CDI (CDI-CORE-2015-505) and the Foundation for Barnes-Jewish Hospital (3770). The Washington University Diabetes Research Center (P30DK020579) provided support for the microscopy. We thank Nicholas White and Shriya Swaminathan for technical assistance.

